# Curvature-sensitive *trans*-assembly of human Atg8-family proteins in autophagy-related membrane tethering

**DOI:** 10.1101/870857

**Authors:** Saki Taniguchi, Masayuki Toyoshima, Tomoyo Takamatsu, Joji Mima

## Abstract

In macroautophagy, de novo formation of the double membrane-bound organelles, termed autophagosomes, is essential for engulfing and sequestering the cytoplasmic contents to be degraded in the lytic compartments such as vacuoles and lysosomes. Atg8-family proteins have been known to be responsible for autophagosome formation via membrane tethering and fusion events of precursor membrane structures. Nevertheless, how Atg8 proteins act directly upon autophagosome formation still remains enigmatic. Here, to further gain molecular insights into Atg8-mediated autophagic membrane dynamics, we study the two representative human Atg8 orthologs, LC3B and GATE-16, by quantitatively evaluating their intrinsic potency to physically tether lipid membranes in a chemically defined reconstitution system using purified Atg8 proteins and synthetic liposomes. Both LC3B and GATE-16 retained the capacities to trigger efficient membrane tethering at the protein-to-lipid molar ratios ranging from 1:100 to 1:5,000. These human Atg8-mediated membrane tethering reactions require *trans*-assembly between the membrane-anchored forms of LC3B and GATE-16 and can be reversibly and strictly controlled by the membrane attachment and detachment cycles. Strikingly, we further uncovered distinct membrane curvature dependences of LC3B- and GATE-16-mediated membrane tethering reactions: LC3B can drive tethering more efficiently than GATE-16 for highly-curved small vesicles (*e.g.* 50 nm in diameter), although GATE-16 turns out to be a more potent tether than LC3B for flatter large vesicles (*e.g.* 200 and 400 nm in diameter). Our findings establish curvature-sensitive *trans*-assembly of human Atg8-family proteins in reconstituted membrane tethering, which recapitulates an essential subreaction of the biogenesis of autophagosomes in vivo.

## Introduction

Macroautophagy, hereafter simply referred to as autophagy, is a fundamental intracellular trafficking pathway vital for the maintenance of homeostasis in all eukaryotic cells, in which the cytoplasmic contents including cytosolic proteins and organelles are selectively or non-selectively delivered to the lytic subcellular compartments, such as vacuoles in yeast and lysosomes in mammalian cells, and then degraded by the lumenal hydrolases [1-6]. During the autophagic processes, eukaryotic cells engulf and sequester the cytoplasmic materials by newly forming the unique double membrane-bound organelles, termed autophagosomes, via expansion and sealing of the small cup-shaped precursor membrane structures, termed phagophores or isolation membranes [1-6]. After sealing of fully expanded phagophore membranes, the completed spherical autophagosomes are eventually transported to and fused with the lytic compartments to release the cargoes into the lumen [1-8]. It is noteworthy that the original 14 autophagy-related (Atg) proteins, which were initially identified in the budding yeast *Saccharomyces cerevisiae* by the pioneering genetic screening experiments of Ohsumi and colleagues [9, 10], are all known to be involved in the biogenesis of autophagosomes [1-8].

Among over 30 Atg protein families that have been identified in budding yeast and higher eukaryotes [1-6], Atg8-family proteins are currently recognized as a conserved key component of the molecular machinery to drive autophagosome formation, in particular, at the phagophore expansion and sealing steps [1-6]. Both yeast Atg8p and the Atg8 orthologs in mammalian cells, including LC3 (microtubule-associated protein 1 light chain 3), GATE-16 (Golgi-associated ATPase enhancer of 16 kDa), and GABARAP (gamma-aminobutyric acid receptor-associated protein), are covalently conjugated to a headgroup of phosphatidylethanolamine (PE) at the C-terminus and thereby specifically localized at autophagosomal membrane compartments through the post-translational lipidation [11-14]. A number of previous cell biological and genetic studies in yeast and mammalian cells demonstrated that the PE-conjugated, membrane-anchored forms of Atg8-family proteins are required for controlling the size and shape of autophagosomes [12-19]. More recently, using a chemically defined system reconstituted with purified Atg8 proteins and synthetic liposomal membranes, it has been further shown that yeast Atg8p and the mammalian Atg8 orthologs have the inherent potency to directly mediate membrane tethering and hemifusion or full fusion by themselves [20-25]. These experimental observations in vivo and in vitro support the idea that Atg8-family proteins function as a membrane tether and fusogen for promoting the tethering and fusion events of expanding phagophores or isolation membranes during autophagosome formation [3-8]. However, it should be noted that the intrinsic tethering and fusogenic activities of Atg8-family proteins are critically dependent on non-physiological conditions employed in those in vitro reconstitution experiments, which include substantially high concentrations of nonlamellar-prone lipids (e.g. PE and di-unsaturated lipid species) and also high Atg8 protein-to-lipid molar ratios [20-25]. Thus, how Atg8-family proteins act directly upon autophagic membrane tethering and fusion processes remains enigmatic and still a matter of debate. In the current study, to further explore the molecular functions of Atg8-family proteins in the biogenesis of autophagosomes, we thoroughly evaluated the intrinsic membrane tethering capacities of the two mammalian Atg8 orthologs, human LC3B and GATE-16 proteins, in the reconstituted proteoliposomal systems mimicking their membrane-bound states in vivo and the physiological lipid bilayers of eukaryotic endomembrane organelles.

## Results and Discussion

The Atg8 protein family in human cells consist of at least seven homologous members, which are classified into the two subfamilies, the LC3 subfamily (LC3A, LC3B, LC3C, and LC3B2) and the GABARAP subfamily (GABARAP, GABARAP-like 1, and GATE-16 that is also named as GABARAP-like 2), whereas only a single Atg8-family protein, Atg8p, is encoded in the budding yeast *S. cerevisiae* (Figure 1A) [21, 25, 26]. All of the human Atg8 orthologs (as well as yeast Atg8p) are a small monomeric globular protein comprised mostly of the Atg8-like domain, exhibiting 30-35% (for the LC3 proteins) and 54-56% (for the GABARAP proteins) sequence identities to Atg8p in yeast (Figure 1A, B; Figure S1) [26]. In the current reconstitution studies testing the intrinsic membrane tethering activities of human Atg8 proteins (Figure 2-6), we selected the two representative Atg8-family members (Figure 1B); LC3B from the LC3 subfamily, which was identified as the first mammalian Atg8 protein that localizes at autophagosomal membrane compartments and plays a role in autophagosome formation [13, 14], and GATE-16 from the GABARAP subfamily, which was initially identified as a novel factor functioning in intra-Golgi membrane trafficking [27-29] and was later reported to be localized at autophagosomal membranes and involved in the biogenesis of autophagosomes [14, 19]. Recombinant LC3B and GATE-16 proteins used in reconstituted liposome tethering assays (Figure 2-6) were purified as the matured, C-terminally truncated forms with an additional artificially-modified polyhistidine tag (His12) following the C-terminal Gly120 residue for LC3B and the Gly116 residue for GATE-16 (denoted as LC3B-His12 and GATE-16-His12, respectively; Figure 1B, C). Purified LC3B-His12 and GATE-16-His12 proteins, which are mostly a monomeric protein (Figure S1), can be stably and specifically attached to membrane surfaces of liposomes through high-affinity binding of a His12 tag to a DOGS-NTA lipid (1,2-dioleoyl-sn-glycero-3-((N-(5-amino-1-carboxypentyl) iminodiacetic acid)-succinyl)) that was included in the lipid composition used for preparing liposomes (Figure 2A). This artificial membrane-anchoring mode mimics the membrane-bound state of native Atg8 proteins, in which, after proteolytic C-terminal truncation by Atg4 cysteine proteases, the C-terminal glycine residues of Atg8s are conjugated to a PE lipid in lipid bilayers of autophagosomal membranes [11, 14]. Furthermore, as earlier reconstitution studies have pointed out the importance of lipid species and compositions in model membrane systems for Atg8-mediated tethering and fusion reactions [22, 25], we employed for reconstitution the physiologically relevant lipid bilayers, roughly mimicking typical lipid compositions of subcellular organelles destined to be membrane sources of autophagosomes in mammalian cells (Figure 2A) [30-33]. The physiologically-mimicking lipid compositions contained synthetic mono-unsaturated phosphatidylcholine (PC), PE, and phosphatidylserine (PS) lipids, a natural phosphatidylinositol (PI) lipid from soybean, and cholesterol from ovine (Figure 2A). Using the experimental conditions above in the present reconstitution systems, we revisited the physiological significance of the intrinsic potency of human Atg8-family proteins to directly mediate membrane tethering (Figure 2-6).

**Figure 1.**
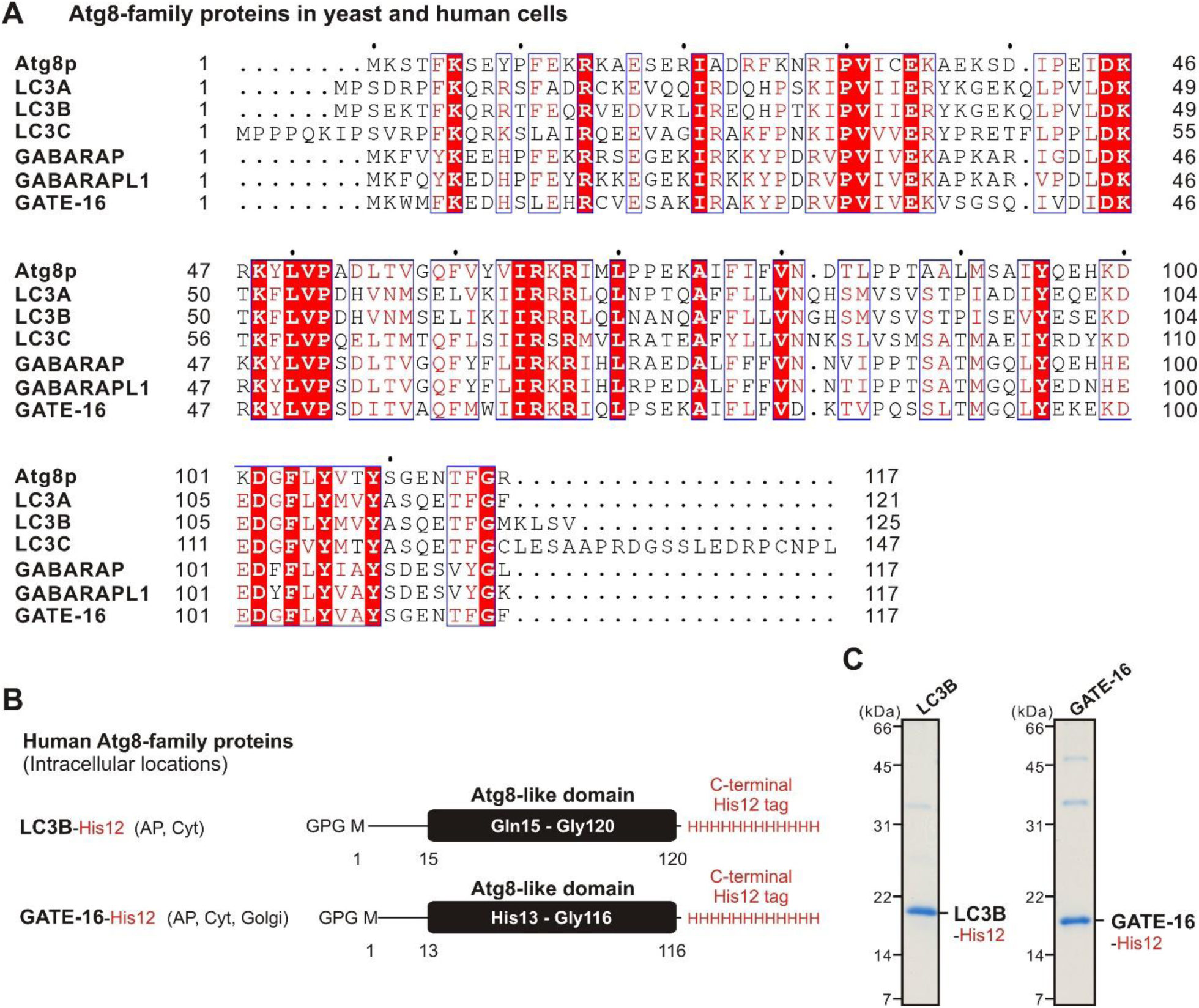
Human Atg8-family proteins used in the current reconstitution studies. (**A**) Sequence alignment of Atg8-family proteins in the yeast *Saccharomyces cerevisiae* (Atg8p) and human cells (LC3A, LC3B, LC3C, GABARAP, GABARAPL1, GATE-16). Amino acid sequences of yeast Atg8p and the human Atg8 orthologs were obtained from UniProtKB (https://www.uniprot.org/), aligned using ClustalW (https://www.genome.jp/tools-bin/clustalw), and rendered with ESPript 3.0 (http://espript.ibcp.fr/ESPript/ESPript/). Identical and similar residues in the alignment are highlighted in red boxes and in red characters, respectively. (**B**) Schematic representation of the C-terminally His12-tagged forms of human Atg8-family proteins used (LC3B-His12, GATE-16-His12), which contain three extra residues (Gly-Pro-Gly) at the N-terminus, the Met1-Gly120 residues for LC3B or the Met1-Gly116 residues for GATE-16, and a polyhistidine tag (His12) at the C-terminus. The intracellular locations of LC3B and GATE-16 indicated include autophagosome (AP), cytosol (Cyt), and Golgi apparatus (Golgi). (**C**) Coomassie blue-stained gels of purified LC3B-His12 and GATE-16-His12 proteins tested in the reconstituted liposome tethering assays in Figures 2-6.

**Figure 2.**
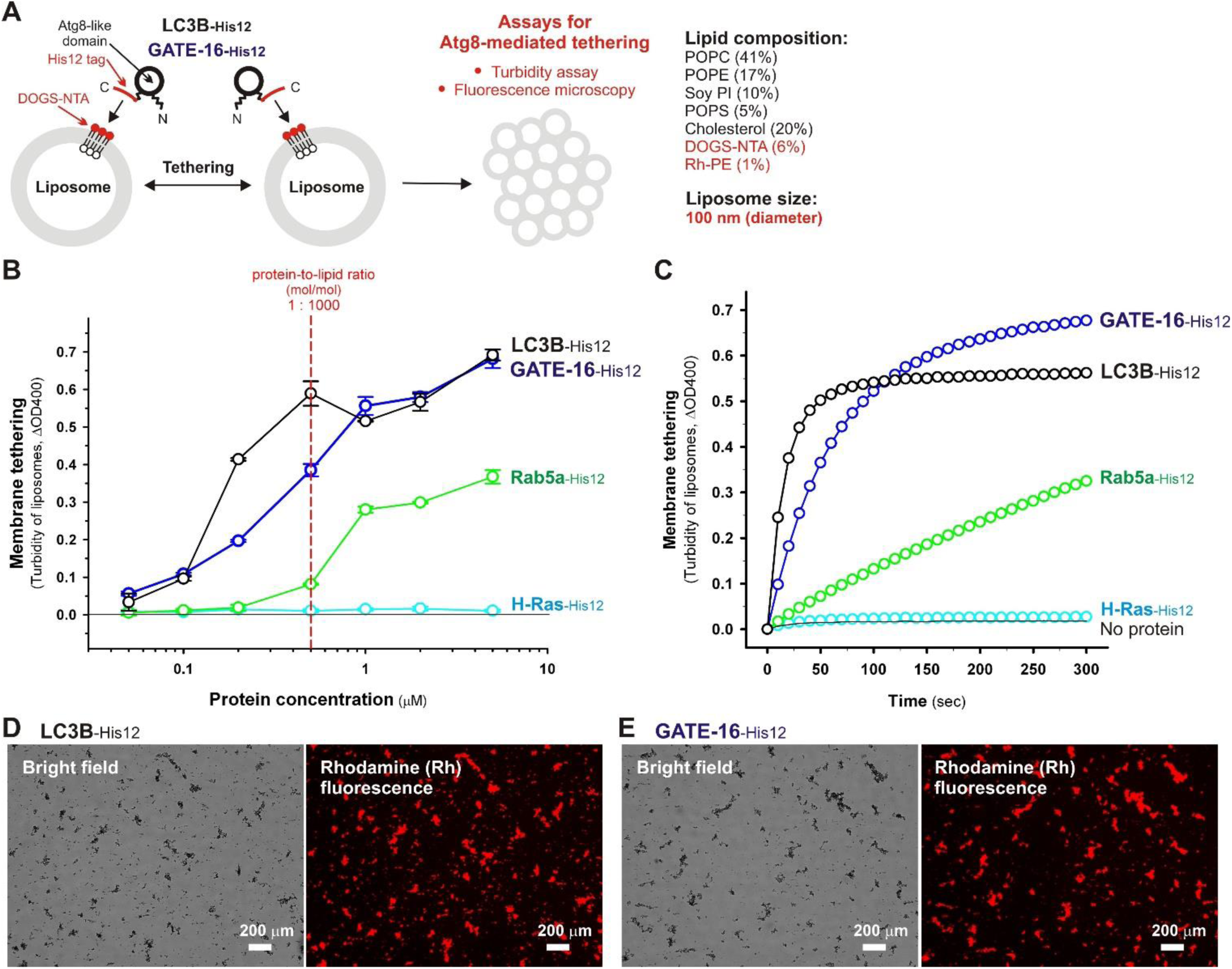
Human Atg8-family proteins, LC3B and GATE-16, can drive efficient and rapid membrane tethering in a chemically defined reconstitution system. (**A**) Schematic representation of reconstituted membrane tethering assays using purified human Atg8-family proteins, LC3B-His12 and GATE-16-His12, and synthetic liposomes bearing a DOGS-NTA lipid. (**B**) End point liposome turbidity assays testing human Atg8-mediated membrane tethering. Purified LC3B-His12, GATE-16-His12, Rab5a-His12, or H-Ras-His12 proteins (0.05, 0.1, 0.2, 0.5, 1, 2, or 5 μM final) were mixed and incubated with DOGS-NTA-bearing liposomes (0.5 mM lipids, 100 nm diameter). After the incubation (30°C, 30 min), turbidity changes of the liposome suspensions were measured with the optical density at 400 nm (ΔOD400). The protein concentration at the protein-to-lipid molar ratio of 1:1,000 (mol/mol) is indicated as a red dashed line. Error bars, S.D. (**C**) Kinetic liposome turbidity assays testing human Atg8-mediated membrane tethering. Turbidity changes of the tethering reactions containing DOGS-NTA-bearing liposomes (1 mM lipids, 100 nm diameter) were monitored in the presence of purified LC3B-His12, GATE-16-His12, Rab5a-His12, or H-Ras-His12 proteins (2 μM final) by measuring ΔOD400 at room temperature for 5 min. (**D, E**) Fluorescence microscopy analysis of liposome clusters induced by human Atg8-mediated membrane tethering. Fluorescence-labeled liposomes bearing rhodamine-PE (Rh-PE) and DOGS-NTA (1 mM lipids, 100 nm diameter) were mixed with purified proteins (2 μM final) of LC3B-His12 (D) or GATE-16-His12 (E), incubated (30°C, 2 h), and analyzed by fluorescence microscopy to obtain the bright field images and Rh-fluorescence images of the liposome tethering reactions. Scale bars, 200 μm.

**Figure 3.**
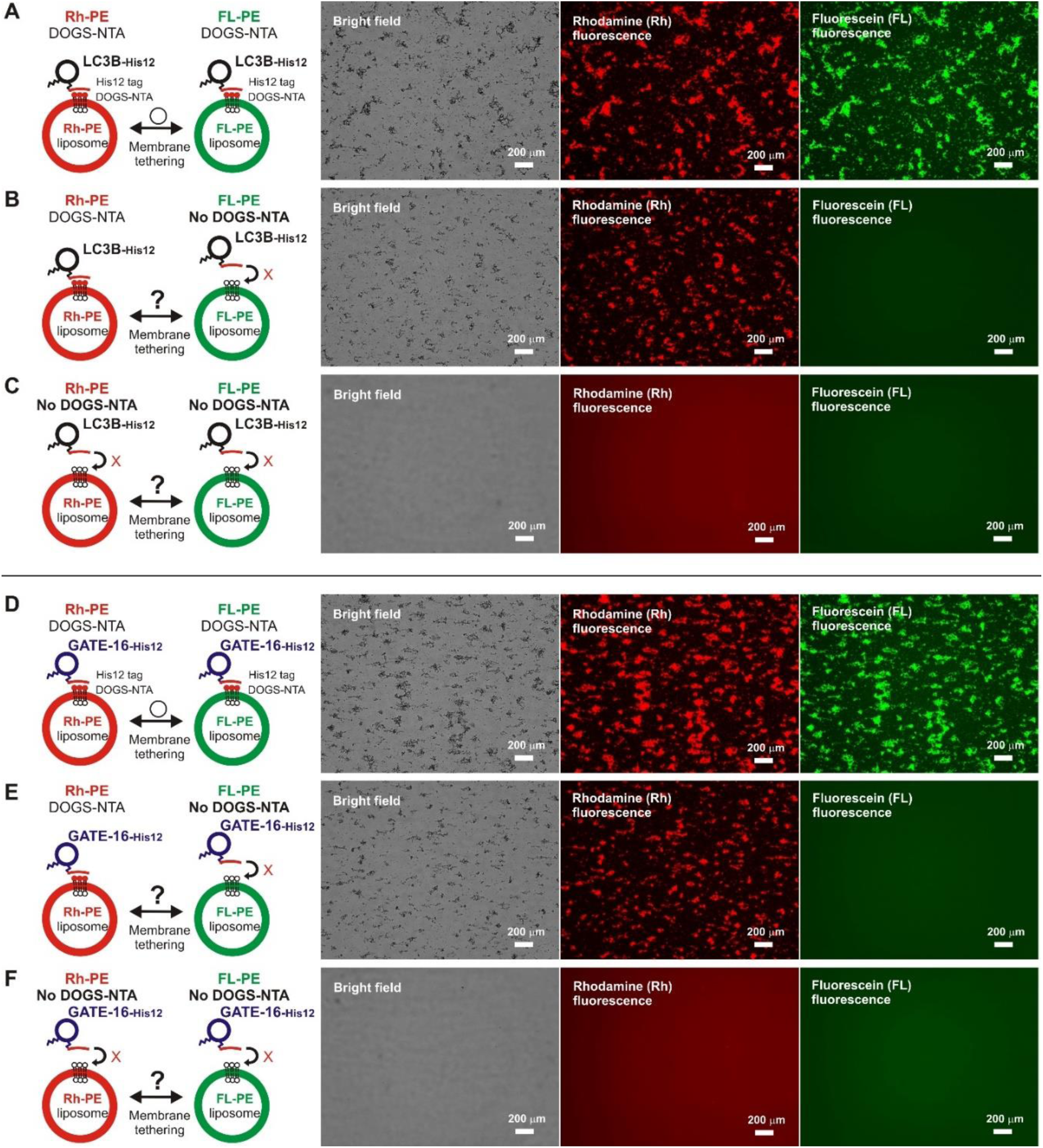
*Trans*-assembly of membrane-anchored forms of human Atg8-family proteins is required for driving reconstituted membrane tethering. (**A-C**) Fluorescence microscopy analysis of reconstituted liposome tethering driven by *trans*-assembly of membrane-anchored LC3B proteins. LC3B-His12 (2 μM) and two types of fluorescence-labeled liposomes, rhodamine-PE (Rh-PE)-bearing liposomes and fluorescein-PE (FL-PE)-bearing liposomes (1 mM lipids for each, 100 nm diameter), were mixed, incubated (30°C, 1 h), and analyzed by fluorescence microscopy to obtain bright field images, Rh-fluorescence images, and FL-fluorescence images of the tethering reactions. DOGS-NTA lipids were present in both the Rh-PE and FL-PE liposomes (A), only in the Rh-PE liposomes (B), or not present in either of the liposomes (C). Scale bars, 200 μm. (**D-F**) Fluorescence microscopy analysis of reconstituted liposome tethering driven by *trans*-assembly of membrane-anchored GATE-16 proteins. GATE-16-His12 and the two types of fluorescence-labeled liposomes were mixed, incubated, and subjected to fluorescence microscopy, as described above for LC3B in (A-C). DOGS-NTA lipids were present in both the Rh-PE and FL-PE liposomes (D), only in the Rh-PE liposomes (E), or not present in either of the liposomes (F). Scale bars, 200 μm.

**Figure 4.**
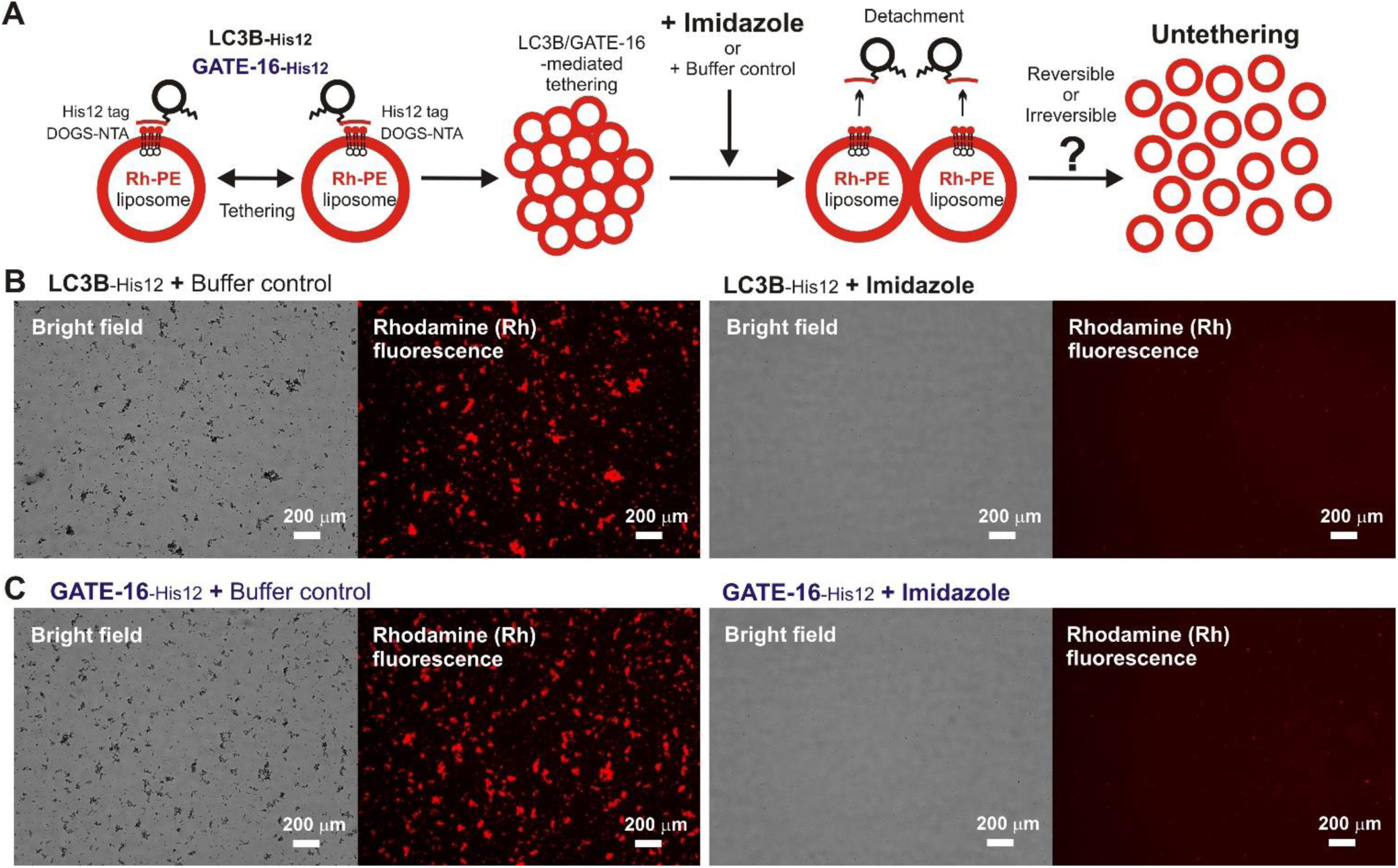
Human Atg8-induced liposome clustering is a non-fusogenic, reversible membrane tethering reaction. (**A**) Schematic representation of fluorescence microscopy analysis for the reversibility of reconstituted membrane tethering mediated by human Atg8-family proteins, LC3B and GATE-16, shown in (B, C). (**B**) The reversibility of reconstituted LC3B-mediated liposome tethering. LC3B-His12 (2 μM final) and rhodamine-PE (Rh-PE)/DOGS-NTA-bearing liposomes (1 mM lipids in final, 100 nm diameter) were mixed and incubated (30°C, 1 h) to induce the formation of LC3B-mediated liposome clusters. After the incubation, the liposome tethering reactions were supplemented with the buffer control or imidazole (250 mM final) to detach LC3B-His12 proteins from DOGS-NTA-bearing liposomes, further incubated (30°C, 1 h), and subjected to fluorescence microscopy. Bright field images and Rh-fluorescence images of the reactions with the buffer control (left panels) or with 250 mM imidazole (right panels) were obtained as in Figure 2D, E. Scale bars, 200 μm. (**C**) The reversibility of reconstituted GATE-16-mediated liposome tethering. GATE-16-His12 and Rh-PE/DOGS-NTA-bearing liposomes were mixed, incubated, supplemented with imidazole or the buffer control, further incubated, and subjected to fluorescence microscopy, as described above for LC3B in (B). Bright field images and Rh-fluorescence images of the GATE-16-containing reactions with the buffer control (left panels) or with imidazole (right panels) were obtained as in Figure 2D, E. Scale bars, 200 μm.

**Figure 5.**
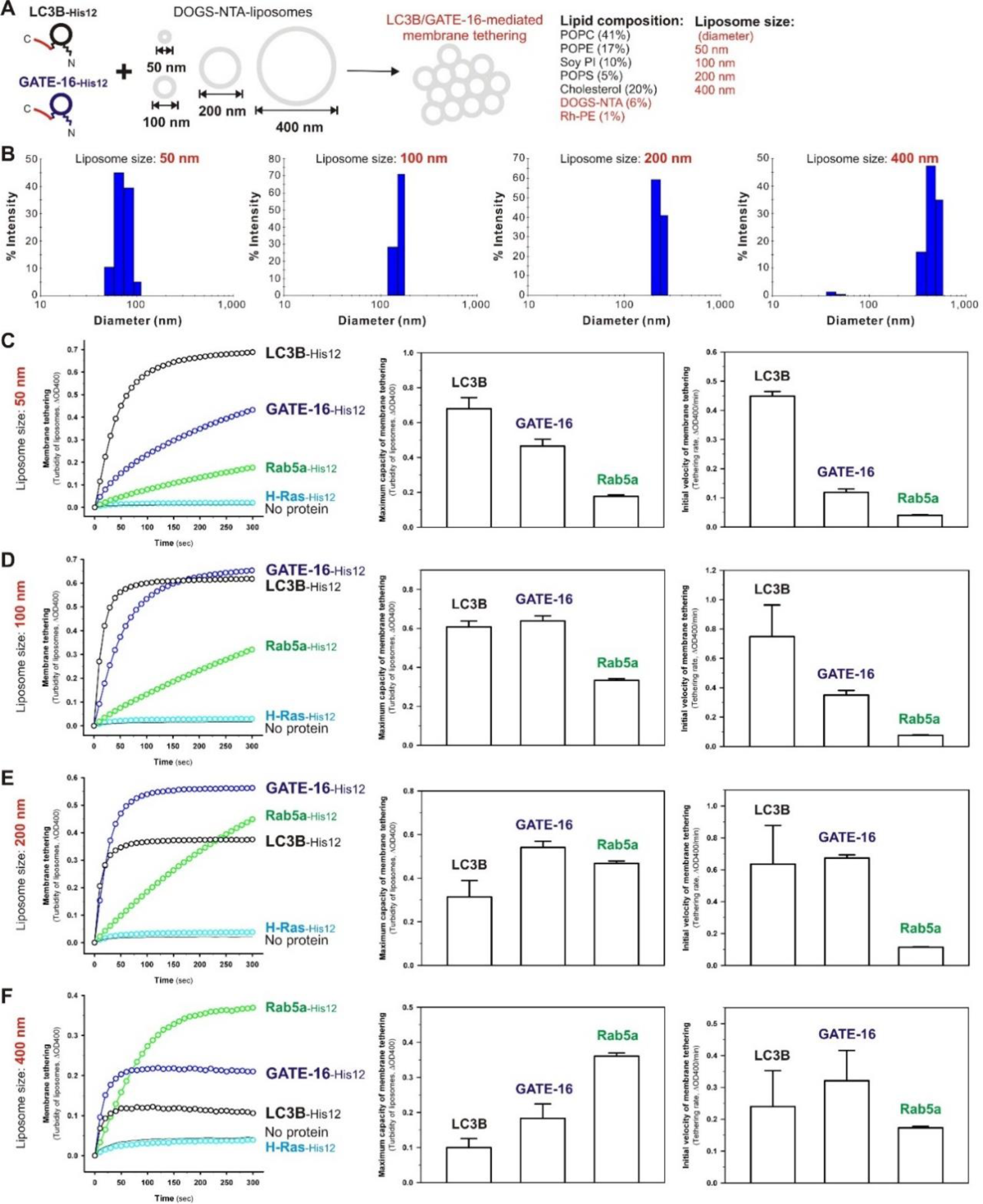
Membrane curvature sensitivity of human Atg8-mediated membrane tethering. (**A**) Schematic representation of reconstituted membrane tethering reactions with human Atg8-family proteins, LC3B-His12 and GATE-16-His12, and four different sizes of DOGS-NTA-bearing liposomes having 50-, 100-, 200-, and 400-nm diameters. (**B**) Dynamic light scattering measurements of the DOGS-NTA-bearing liposomes used, showing histograms of size distributions of the 50-, 100-, 200-, and 400-nm-diameter liposomes. (**C-F**) Kinetic liposome turbidity assays testing membrane curvature dependences of LC3B- and GATE-16-mediated membrane tethering reactions. LC3B-His12, GATE-16-His12, Rab5a-His12, or H-Ras-His12 proteins (2 μM in final) and the DOGS-NTA-bearing 50-nm (C), 100-nm (D), 200-nm (E), or 400-nm (F) liposomes (1 mM lipids in final) were mixed and then immediately assayed for turbidity changes by measuring ΔOD400, as in Figure 2C. The kinetic turbidity data obtained (left panels) were further quantitatively analyzed by curve fitting to determine means and S.D. values of the maximum tethering capacities (middle panels) and the initial tethering velocities (right panels). The maximum tethering capacities of LC3B for 50-nm (C), 200-nm (E), and 400-nm (F) liposomes are significantly different from those of GATE-16, and the initial tethering velocities of LC3B for 50-nm (C) and 100-nm (D) liposomes are significantly different from those of GATE-16 (*p* < 0.05, calculated using one-way ANOVA). Error bars, S.D.

**Figure 6.**
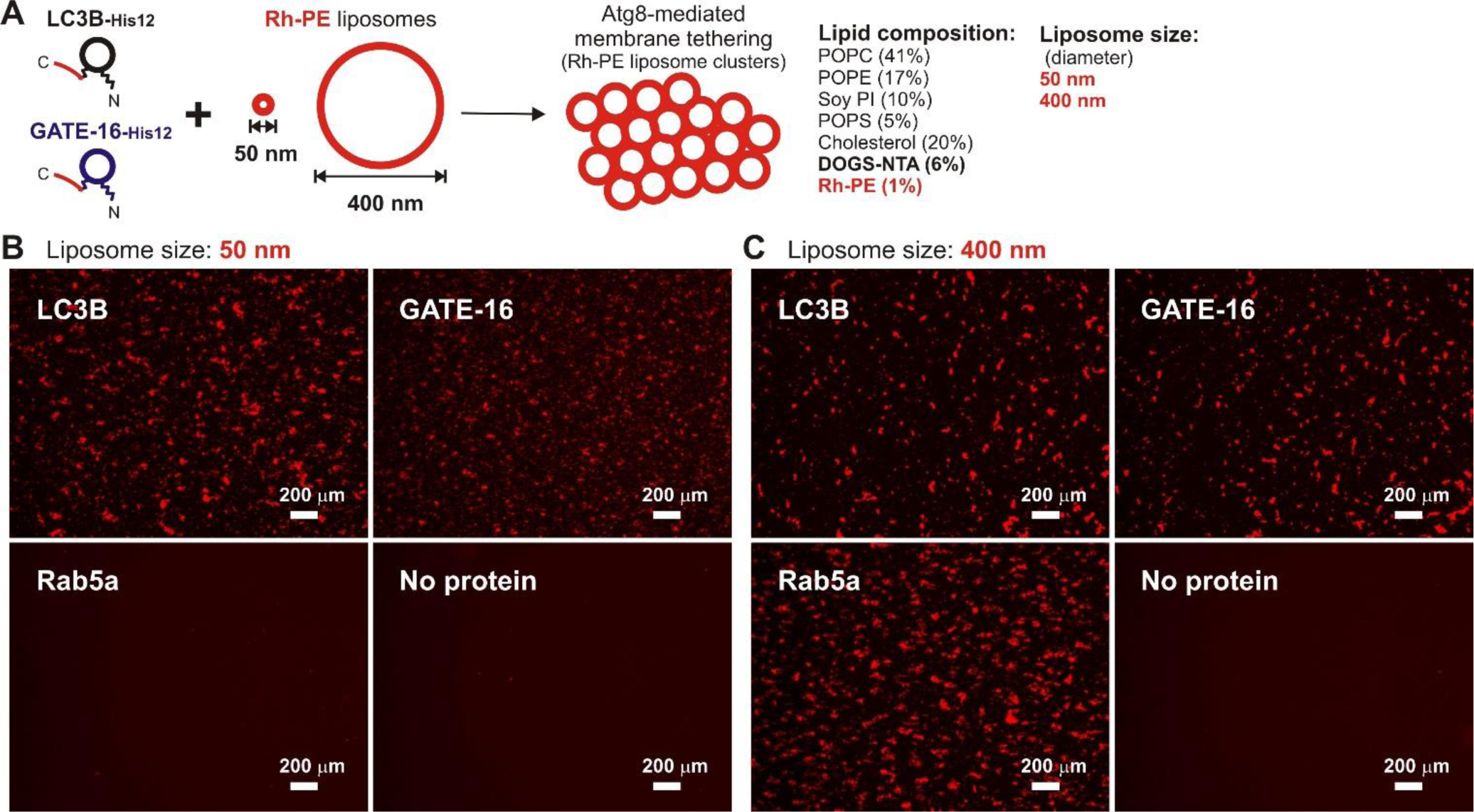
Efficient clustering of highly-curved liposomes induced by human Atg8-mediated membrane tethering. (**A**) Schematic representation of fluorescence microscopy analysis of human Atg8-mediated membrane tethering of small highly-curved 50-nm liposomes and larger-sized flatter 400-nm liposomes in (B, C). (**B, C**) Rhodamine (Rh) fluorescence images of liposome clusters induced by human Atg8-mediated tethering of the 50-nm liposomes (B) and 400-nm liposomes (C). Rh-PE/DOGS-NTA-bearing 50-nm (B) and 400-nm (C) liposomes (1 mM lipids) were mixed with LC3B-His12, GATE-16-His12, or Rab5a-His12 (2 μM final), incubated (30°C, 1 h), and subjected to fluorescence microscopy to obtain Rh-fluorescence images. Scale bars, 200 μm.

### *Trans*-assembly of human Atg8 proteins can drive membrane tethering in a physiological context

To comprehensively and quantitatively evaluate the intrinsic membrane tethering capacities of human LC3B and GATE-16 proteins in a chemically defined proteoliposomal system, we employed three types of reconstituted liposome tethering assays, which had been previously developed to test membrane tethering driven by human Rab-family small GTPases in our recent studies [34-37], including a high-throughput 384-well microplate-based endpoint assay to measure the turbidity of liposomes (Figure 2B), a kinetic assay to monitor liposome turbidity changes (Figure 2C), and a fluorescence cell counter-based imaging assay giving an unbiased quantitative data of clusters of tethered liposomes (Figure 2D, E). Strikingly, both of the human Atg8 proteins exhibited the abilities to drive highly efficient and rapid membrane tethering by themselves, in the absence of any other additional protein components (Figure 2B-E). LC3B-His12 and GATE-16-His12 proteins triggered efficient tethering of synthetic, but physiologically-mimicking, DOGS-NTA-bearing liposomes (100 nm in diameter; Figure 2A) even at the protein-to-lipid molar ratios of 1:1,000 or below (black and blue circles, respectively, in Figure 2B). However, Rab5a-His12, which has been so far found to be the most active membrane tether among human Rab-family small GTPase isoforms [34-37], had the substantially less tethering capacity under the same conditions (green circles, Figure 2B) and, H-Ras-His12, used as the negative control of a C-terminally His12-tagged protein, was completely unable to initiate liposome tethering in the turbidity assays (cyan circles, Figure 2B). Moreover, the kinetic liposome turbidity assays indicated that the initial velocities of human LC3B- and GATE-16-mediated membrane tethering reactions were much higher than that of the Rab5a-mediated reaction, when tested at the protein-to-lipid ratios of 1:500 (Figure 2C). Consistent with those data from the turbidity assays (Figure 2B, C), LC3B-His12 and GATE-16-His12 were able to induce the formation of massive clusters of the liposomes (100 nm in diameter) bearing DOGS-NTA and rhodamine-PE (Rh-PE) through their intrinsic tethering activities (Figure 2D, E). In the fluorescence imaging assays employed at the protein-to-lipid ratios of 1:500 (Figure 2D, E), LC3B- and GATE-16-induced liposome clusters had the average particle sizes of 420 μm^2^ and 680 μm^2^, respectively, whereas an untethered single 100-nm-diameter liposome gives only 0.0079 μm^2^ for its maximum particle size in theory. Thus, all of the experimental data from these three independent reconstitution assays of liposome tethering faithfully reflect the very high efficiency of LC3B- and GATE-16-mediated membrane tethering reactions.

In the endpoint turbidity assays, we tested the intrinsic membrane tethering potency of LC3B and GATE-16 proteins comprehensively at the protein-to-lipid molar ratios ranging from 1:10,000 to 1:100 (mol/mol), demonstrating that both of the human Atg8-family proteins can trigger membrane tethering at the protein-to-lipid ratio of 1:5,000 and they drive membrane tethering more efficiently at the higher ratios up to 1:100 (black and blue circles, Figure 2B). Since it has been described that protein densities of fusogens or tethers on membrane surfaces are critical to establishing the physiologically-relevant reconstitution systems for protein-driven membrane tethering and fusion mediated by SNARE-family proteins [38-41] and Rab-family small GTPases [35-37], we therefore assessed whether the Atg8 protein densities tested in the current tethering assays (Figure 2B) are relevant to the physiological environments of native Atg8-family proteins associated with autophagic membrane compartments. Previous works by Klionsky and colleagues quantitatively estimated the protein stoichiometry of several representative Atg proteins at the phagophore assembly site (PAS) by a fluorescence microscopic analysis [42] and also the size of the fully expanded phagophore by an electron microscopic analysis of autophagic bodies in the Pep4p-deficient yeast vacuoles [43]. These quantitative analyses gave approximately 272 ± 9 Atg8p molecules at the PAS (mean ± S.E.M., n = 100) [42, 43] and 127 ± 2 nm for the radii of autophagic bodies (mean ± S.E.M., n > 200) [43]. Using the estimated values [42, 43] and the average surface area of the headgroups of phospholipids (0.65 nm^2^) [44], we can calculate the physiological protein-to-lipid ratio for Atg8p on the phagophore as about 1:2,300 (272 Atg8p molecules to 623,000 lipids). Notably, when tested at the protein-to-lipid ratio of 1:2,500 (0.2 μM LC3B/GATE-16 to 0.5 mM lipids; Figure 2B) that is very close to the ratio calculated above, both LC3B and GATE-16 were still able to exhibit the significant intrinsic tethering activities (Figure 2B). In addition, as membrane-anchored Atg8 proteins can be selectively sorted into a highly curved membrane segment by several folds [45, 46], it is conceivable that the protein-to-lipid ratio of 1:500, which was used in the kinetic turbidity assays and fluorescence microscopy assays with 100-nm-diameter liposomes (Figure 2C-E), can provide the physiologically-relevant Atg8 density on membrane surfaces. It should be also noted that, assuming that Atg8 proteins are a typically spherical 15-kDa protein with an average radius of 1.6 nm [47], membrane-anchored LC3B and GATE-16 proteins only occupy approximately 4.5% of the surface areas of liposomes in the current reconstitution systems, when tested at the protein-to-lipid ratio of 1:500 (Figure 2).

Next, we asked whether human Atg8-mediated membrane tethering is primarily driven by *trans*-assembly between membrane-anchored Atg8 proteins on two distinct lipid bilayers (Figure 3). Although the prior study for yeast Atg8p demonstrated the presence of the dimers and oligomers in the reconstituted tethering and fusion reactions [20], it remains unclear whether Atg8 proteins assemble into the homo-complexes in *cis* on one membrane or in *trans* on two opposing membranes to be tethered. To address this, we employed the fluorescence imaging assays for LC3B (Figure 3A-C) and GATE-16 (Figure 3D-F) with two types of the fluorescence-labeled DOGS-NTA-liposomes bearing either Rh-PE or fluorescein-PE (FL-PE). LC3B and GATE-16 were able to induce the formation of massive liposome clusters which contain both of the Rh-PE-liposomes (middle panels, Figure 3A, D) and FL-PE-liposomes (right panels, Figure 3A, D). This is consistent with the preceding results in the fluorescence imaging assays using the Rh-PE-liposomes alone (Figure 2D, E). However, when a DOGS-NTA lipid was omitted from the FL-PE-liposomes, these two human Atg8-family proteins no longer had the potency to trigger clustering of the FL-labeled liposomes lacking DOGS-NTA (Figure 3B, E, right panels), whereas they still retained the intrinsic capacities to efficiently and selectively tether the DOGS-NTA-bearing Rh-PE-liposomes in the same tethering reactions (Figure 3B, E, middle panels). Moreover, when both of the Rh-PE-liposomes and FL-PE-liposomes lacked a DOGS-NTA lipid, LC3B and GATE-16 completely lost their membrane tethering activities, giving little or no detectable liposome clusters in the reactions (Figure 3C, F). These results faithfully reflect that LC3B and GATE-16 proteins need to be anchored on both, not either one, of two opposing membrane surfaces for initiating membrane tethering in the current chemically defined systems. Thus, this leads us to conclude that *trans*-assembly of membrane-anchored Atg8 proteins is certainly required for human Atg8-mediated membrane tethering. Additionally, it is noteworthy that the membrane-anchored forms of Atg8-family proteins can specifically associate with membrane-bound Atg8 proteins, but not with the membrane-unbound soluble forms, during membrane tethering, as homo-oligomerization of human LC3B proteins was not observed in GST pull-down assays for LC3B in solution (Figure S2) and liposome co-sedimentation assays testing the interactions of membrane-bound LC3B-His12 proteins with soluble untagged LC3B proteins lacking a His12 tag (Figure S3).

Our current findings so far established that *trans*-assembly of human Atg8-family proteins directly drives efficient and rapid membrane tethering under physiologically-relevant conditions of the Atg8 protein densities on lipid bilayer surfaces (Figure 2, 3). Based on that, next we revisited the question of whether Atg8-mediated membrane tethering is a reversible tethering-only reaction or an irreversible reaction that spontaneously proceeds to hemifusion or fusion between two distinct membranes tethered. To test reversibility of human Atg8-mediated membrane tethering, LC3B- and GATE-16-induced liposome clusters pre-formed after incubation (30°C, 1 h) were supplemented with imidazole that leads to dissociation of LC3B-His12 and GATE-16-His12 proteins from liposomes (Figure 4A), further incubated (30°C, 1 h), and eventually analyzed by the fluorescence imaging assays (Figure 4). Strikingly, LC3B-and GATE-16-mediated liposome clusters were completely or thoroughly disassembled into an undetectable small particle (Figure 4B, C, right panels), although LC3B and GATE-16 still efficiently mediated liposome clustering in the equivalent reactions supplemented with the buffer control instead (Figure 4B, C, left panels). These results establish that human Atg8-mediated membrane tethering reactions are a reversible process that can be strictly controlled by the membrane attachment and detachment cycles of Atg8 proteins. Furthermore, they strongly suggest that membrane-anchored Atg8 proteins can trigger only a reversible membrane tethering reaction by themselves but not mediate an irreversible hemifusion or fusion events when any other protein components are not present. Additional fusogenic proteins acting together with Atg8 proteins may include autophagy-related SNARE-family proteins that have been reported to be targeted to autophagic compartments and involved in autophagosome formation and maturation [22, 48-50]. However, it should be noted that, since membrane attachment of cytosolic Atg8 proteins via the conjugation to a nonlamellar-prone PE lipid is proposed to take place at locally-curved unstable lipid bilayers [45, 46, 51], we will not exclude the possibility that the intrinsic fusogenic potency of membrane-anchored Atg8 proteins directly facilitate membrane fusion events specifically at the highly-curved, fusion-prone membrane segments of autophagic compartments, including the rim of the cup-shaped phagophore membrane.

### Curvature sensitivity of human LC3B- and GATE-16-mediated membrane tethering

During autophagosome formation, key protein components in autophagy specifically associate with and function at the precursor membrane structures exhibiting a variety of the membrane surface curvatures, such as Golgi-derived Atg9-containing small vesicles, tubular protrusions at the ER (endoplasmic reticulum) and recycling endosomes for the membrane sources of autophagosomes, and the growing phagophore membranes having the strongly-curved rim and relatively-flatter concave inner and convex outer surfaces [46]. Atg8-family proteins in yeast and mammalian cells, indeed, have been reported to localize at the phagophore rims and also both sides of the double-membrane organelles after the PE-conjugation for playing their multiple roles in autophagosome formation and maturation [7, 8, 12-14, 51]. Therefore, we finally asked whether membrane curvature is critical for human Atg8-mediated membrane tethering driven by *trans*-assembly of membrane-anchored Atg8 proteins, by quantitatively investigating the intrinsic tethering activities of LC3B and GATE-16 for liposomal membranes having different particle sizes (Figure 5). Four types of DOGS-NTA-bearing liposomes were prepared by extrusion through 50-, 100-, 200-, and 400-nm pore sizes (Figure 5A) and evaluated for the size distributions using dynamic light scattering, establishing that the liposome preparations had the mean diameters roughly similar to the theoretical pore sizes used and that their size distributions were significantly different from each other (Figure 5B). Using these 50-, 100-, 200-, and 400-nm-diameter liposome preparations (Figure 5A, B), we employed the kinetic turbidity assays for LC3B, GATE-16, and Rab5a that is a putative membrane tether functioning in non-autophagic, endocytic trafficking pathways [34-37, 52], followed by determining their maximum tethering capacities and initial tethering velocities from curve fitting of the kinetic data obtained (Figure 5C-F).

Strikingly, LC3B and GATE-16, the autophagic membrane tethers, triggered very efficient and rapid tethering of the 50-nm- and 100-nm-diameter liposomes (black and blue circles, Figure 5C, D), while the endocytic membrane tether Rab5a was substantially less competent to drive tethering of these small highly-curved liposomes (green circles, Figure 5C, D). In contrast, Rab5a retained the capacity to cause membrane tethering for the larger-sized, flatter liposomes (e.g. 400 nm in diameter) more efficiently than LC3B and GATE-16 (see the maximum tethering capacities, middle panel, Figure 5F), even though the two autophagic tethers were still able to initiate rapid membrane tethering of the large lipid vesicles (see the initial tethering velocities, right panel, Figure 5F). These membrane curvature dependences of the autophagic and endocytic membrane tethers in the kinetic turbidity assays are fully consistent with the results from the fluorescence imaging assays, which were employed using the 50-nm- and 400-nm-diameter liposomes (Figure 6). The obtained fluorescence images clearly indicated that only the two autophagic tethers LC3B and GATE-16 (but not Rab5a) had the intrinsic tethering activities to induce the formation of massive clusters of the small highly-curved 50-nm liposomes (Figure 6B). However, for the large flatter 400-nm liposomes, Rab5a-mediated membrane tethering was significantly more efficient than that mediated by the autophagic tethers (Figure 6C). Furthermore, it is quite Intriguing that we uncover distinct membrane curvature dependences of the two different autophagic membrane tethers, LC3B and GATE-16, in reconstituted membrane tethering (Figure 5C-F). In the kinetic turbidity assays with the smallest, highly-curved 50-nm-diameter liposomes (Figure 5C), LC3B can specifically function as a hyperactive membrane tether, in comparison to the other autophagic tether GATE-16, yielding its initial tethering velocity almost 4-fold higher than that of GATE-16 (see the initial tethering velocities, right panel, Figure 5C). Nevertheless, when tested the same kinetic turbidity assays but with the 200-nm and 400-nm liposomes (Figure 5E, F), GATE-16 turned out to be a significantly more potent membrane tether, rather than LC3B, for the larger-sized flatter lipid vesicles (see the maximum tethering capacities, middle panels, Figure 5E, F). It should also be noted that, even though there were no statistically significant differences between the initial tethering velocities of LC3B and GATE-16 for the large 200-m/400-nm liposomes, LC3B exhibited the substantially lower tethering capacities than those of GATE-16 under the conditions (Figure 5E, F). These data lead to speculation that *cis*-interactions between LC3B proteins anchored to the same membrane can occur on a relatively flat lipid bilayer, thereby preventing the stable protein assemblies in *trans* on two opposing membranes.

Our novel findings on the curvature sensitivity of human LC3B- and GATE-16-mediated membrane tethering lead us to postulate that the specific molecular functions of diverse Atg8-family members in human and mammalian cells rely, at least in part, upon their distinct curvature dependences for the intrinsic membrane tethering potency. Since cytological studies on mammalian Atg8 orthologs previously reported that the LC3-subfamily members appear to be involved in the phagophore elongation step whereas the GATE-16/GABARAP subfamily proteins is found to be critical for the later stages such as autophagosome closure and maturation [19], the curvature sensitivity of membrane tethering may enable the LC3 and GATE-16/GABARAP subfamilies to specifically act at the different stages during the biogenesis of autophagosomes. In addition to the working model above, considering that autophagy involves miscellaneous membrane sources having various sizes and shapes for autophagosome formation [46], it is also conceivable that, through the distinct curvature sensitivities, a variety of mammalian Atg8-family members can function as a selective membrane tether exclusively for their cognate membrane sources. Future studies on Atg8-mediated membrane tethering (perhaps, and membrane fusion) using a chemically defined reconstitution system will provide further important insights into the mechanistic details of the selective *trans*-assembly of Atg8 proteins for driving autophagy-related membrane tethering and also into the biological significance of the curvature-sensitive membrane tethering processes mediated by diverse members of the Atg8 protein family.

## Materials and Methods

### Protein expression and purification

The expression vectors for the two human Atg8 orthologs, LC3B (UniProtKB ID: Q9GZQ8) and GATE-16 (UniProtKB ID: P60520), were constructed by the ligation-independent cloning method using a pET-41 Ek/LIC vector kit (Novagen). The coding sequences of human LC3B and GATE-16 proteins were amplified by PCR with KOD-Plus-Neo DNA polymerase (Toyobo), Human Universal QUICK-Clone cDNA II (Clontech) for a template cDNA, and the oligonucleotide primers that were designed to generate the DNA fragments containing the additional sequences encoding a human rhinovirus (HRV) 3C protease site (Leu-Glu-Val-Leu-Phe-Gln-Gly-Pro) upstream of the initial ATG codons and also those encoding polyhistidine residues (His12) downstream of the codons for the Gly120 residue of LC3B and the Gly116 residue of GATE-16 (Figure 1A, 1B). The PCR fragments generated were cloned into a pET-41 Ek/LIC vector (Novagen) expressing an N-terminally GST-His6-tagged protein. The GST-His6-tagged forms of LC3B-His12 and GATE-16-His12 proteins were expressed in *Escherichia coli* BL21(DE3) cells (Novagen) harboring the pET-41-based vectors constructed in Lysogeny Broth (LB) medium (1 liter each) containing kanamycin (final 50 μg/ml). After inducing protein expression by adding IPTG (0.4 mM final, 37°C, 3 h), cultured cells were harvested by centrifugation and resuspended in RB150 buffer (20 mM Hepes-NaOH, pH 7.4, 150 mM NaCl, 10% glycerol) containing 5 mM MgCl2, 1 mM dithiothreitol (DTT), 1 mM phenylmethylsulfonyl fluoride (PMSF), and 1.0 μg/ml pepstatin A (40 ml each). Cell suspensions were freeze-thawed in a liquid nitrogen bath and then a water bath at 30°C, lysed by sonication (UD-201 ultrasonic disrupter, Tomy Seiko), and centrifuged at 50,000 rpm (70 min, 4°C) with a 70 Ti rotor (Beckman Coulter). GST-His6-tagged LC3B-His12 and GATE-16-His12 proteins in the supernatants obtained were isolated by mixing with COSMOGEL GST-Accept beads (50% slurry, 4 ml each; Nacalai Tesque), followed by incubation with gentle agitation (4°C, 3 h). The protein-bound GST-Accept beads were then washed by RB150 containing 5 mM MgCl2 and 1 mM DTT (8 ml each) three times, resuspended in the same buffer (4 ml each) containing HRV 3C protease (32 units each; Novagen), and incubated without agitation (4°C, 16 h). Purified tag-less LC3B-His12 and GATE-16-His12 proteins, which contain the Met1-Gly120 residues for LC3B or the Met1-Gly116 residues for GATE-16 and only three extra residues (Gly-Pro-Gly) at their N-terminus (Figure 1A, 1B), were eluted from the beads by HRV 3C protease cleavage. After centrifugation of the bead suspensions (15,300 x *g*, 10 min, 4°C), protein concentrations of purified LC3B-His12 and GATE-16-His12 proteins in the supernatants obtained (around 4 ml each) were determined using Protein Assay CBB Solution (Nacalai Tesque) and bovine serum albumin (BSA) for a standard protein. Recombinant proteins of human Rab5a-His12 and H-Ras-His12, which were used as a control protein in the reconstituted liposome tethering assays, were expressed in *E. coli* cells and purified as described previously [34, 35, 37].

### Liposome preparation

Non-fluorescent lipids, which include POPC (1-palmitoyl-2-oleoyl-PC), POPE (1-palmitoyl-2-oleoyl-PE), soy PI, POPS (1-palmitoyl-2-oleoyl-PS), DOGS-NTA, and ovine cholesterol, were all purchased from Avanti Polar Lipids. Fluorescence-labeled lipids, Rh-PE and FL-PE, were from Invitrogen. Lipid mixes for preparation of synthetic protein-free liposomes contained POPC [41% (mol/mol)], POPE (17%), soy PI (10%), POPS (5%), cholesterol (20%), DOGS-NTA (6%), and Rh-PE (1%) or FL-PE (1%) in chloroform. Dried lipid films were prepared by removing chloroform from the lipid mixes with a stream of nitrogen gas, resuspended in RB150 containing 5 mM MgCl2 and 1 mM DTT by vortexing, incubated with gentle agitation (37°C, 1 h), and freeze-thawed in liquid nitrogen and a water bath at 30°C. Lipid suspensions obtained (final 8 mM lipids) were extruded 25 times through polycarbonate filters (pore diameters of 50, 100, 200, or 400 nm) in a mini-extruder (Avanti Polar Lipids) at 40°C. Size distributions of the extruded liposomes were measured by dynamic light scattering (DLS) using a DynaPro NanoStar DLS instrument (Wyatt Technology).

### Liposome turbidity assay

The intrinsic membrane tethering activities of human Atg8-family proteins, LC3B and GATE-16, were assayed by measuring turbidity of liposome suspensions in the presence of purified Atg8 proteins, as described previously for the assays of human Rab small GTPase-mediated membrane tethering [34-37]. In the endpoint turbidity assays, purified LC3B-His12 or GATE-16-His12 proteins (0.05-5 μM in final) and the DOGS-NTA-bearing protein-free liposomes (100 nm diameter; final 0.5 mM total lipids), both of which had been separately incubated in RB150 containing 5 mM MgCl2 and 1 mM DTT (30°C, 10 min), were mixed (total 150 μl) and transferred into a black 384-well plate with a clear flat bottom (40 μl per well; Corning 3544), followed by incubation without agitation (30°C, 30 min). After the 30-min incubation, turbidity of the liposome suspensions in a 384-well plate was measured with optical density at 400 nm (OD400) on a SpectraMax Paradigm plate reader (Molecular Devices), as described [36, 37]. For a control, the C-terminally His12-tagged non-Atg8-family proteins, human Rab5a-His12 and H-Ras-His12, were also incubated with protein-free liposomes in a 384-well plate and assayed for turbidity changes, as for the Atg8-containing turbidity reactions. All of the turbidity data (ΔOD400) were corrected by subtracting the values of the liposome-only reactions without any protein components. Means and standard deviations of the ΔOD400 data were determined from three independent experiments (Figure 2B).

Kinetic assays for Atg8-mediated membrane tethering were employed by measuring turbidity of liposome suspensions, as previously described for Rab GTPase-mediated liposome tethering [34-37]. After separately preincubating purified LC3B-His12 or GATE-16-His12 proteins (2 μM in final) and the DOGS-NTA-bearing liposomes (50, 100, 200, or 400 nm in diameter; final 1 mM lipids) in RB150 containing 5 mM MgCl2 and 1 mM DTT (30°C, 10 min), they were mixed (total 160 μl), applied into a 10-mm path-length cell (105.201-QS, Hellma Analytics) in a DU720 spectrophotometer (Beckman Coulter), and then immediately measured with monitoring the optical density changes at 400 nm (ΔOD400) at room temperature (5 min, 10-sec intervals). In addition, Rab5a-His12 and H-Ras-His12 proteins were also assayed as above for LC3B and GATE-16. To quantitatively analyze the kinetics of the Atg8-mediated liposome tethering reactions, the turbidity data obtained were subjected to curve fitting using ImageJ2 software (National Institutes of Health) and the logistic function formula, y = a/(1+b*exp(-c*x)), in which y and x are the ΔOD400 value and the time (min), respectively. The maximum capacities of membrane tethering were defined as the theoretical maximum ΔOD400 values of the fitted curves at t = ∞, which can be calculated as “a” from the formula above. The initial velocities were defined as the maximum slopes of the fitted curves, which can be calculated as “c*a/4” from the formula above. Means and standard deviations of the maximum capacities and initial velocities of membrane tethering were determined from three independent experiments. The maximum tethering capacities and the initial tethering velocities were statistically evaluated using one-way ANOVA in SigmaPlot 11 (Systat Software). Statistical differences among data sets were considered significant at *p* < 0.05. All of the kinetic assay data shown were obtained from one experiment and were typical of those from more than three independent experiments.

### Fluorescence microscopy

Fluorescence microscopy assays for the liposome clusters induced by human Atg8-mediated membrane tethering were employed using a LUNA-FL automated fluorescence cell counter (Logos Biosystems) and LUNA cell counting slides (L12001, Logos Biosystems), as described for human Rab small GTPase-mediated membrane tethering [36, 37]. Fluorescence-labeled liposomes bearing Rh-PE or FL-PE (50, 100, or 400 nm diameter; final 1 or 2 mM lipids) and purified LC3B-His12, GATE-16-His12, or Rab-His12 proteins (final 2 μM) were preincubated separately (30°C, 10 min), mixed in RB150 containing 5 mM MgCl2 and 1 mM DTT (total 80 μl), further incubated (30°C, 60 min), and then applied to a well of the cell counting slides (15 μl per well). Bright field images, Rh-fluorescence images, and FL-fluorescence images of the liposome tethering reactions were obtained and processed using the LUNA-FL fluorescence cell counter. Particle sizes of liposome clusters in the fluorescence images were analyzed using the ImageJ2 software with setting the lower intensity threshold level to 150, the upper intensity threshold level to 255, and the minimum particle size to 10 pixel^2^, as described [35-37].

## Acknowledgements

We thank Dr. Genji Kurisu and Dr. Hideaki Tanaka (Institute for Protein Research, Osaka University) for access to dynamic light scattering experiments. This study was supported by Grants-in-Aid for Scientific Research from the Ministry of Education, Culture, Sports, Science and Technology, Japan (MEXT) (to J.M.).

## Supplemental Figures

**Figure S1.**
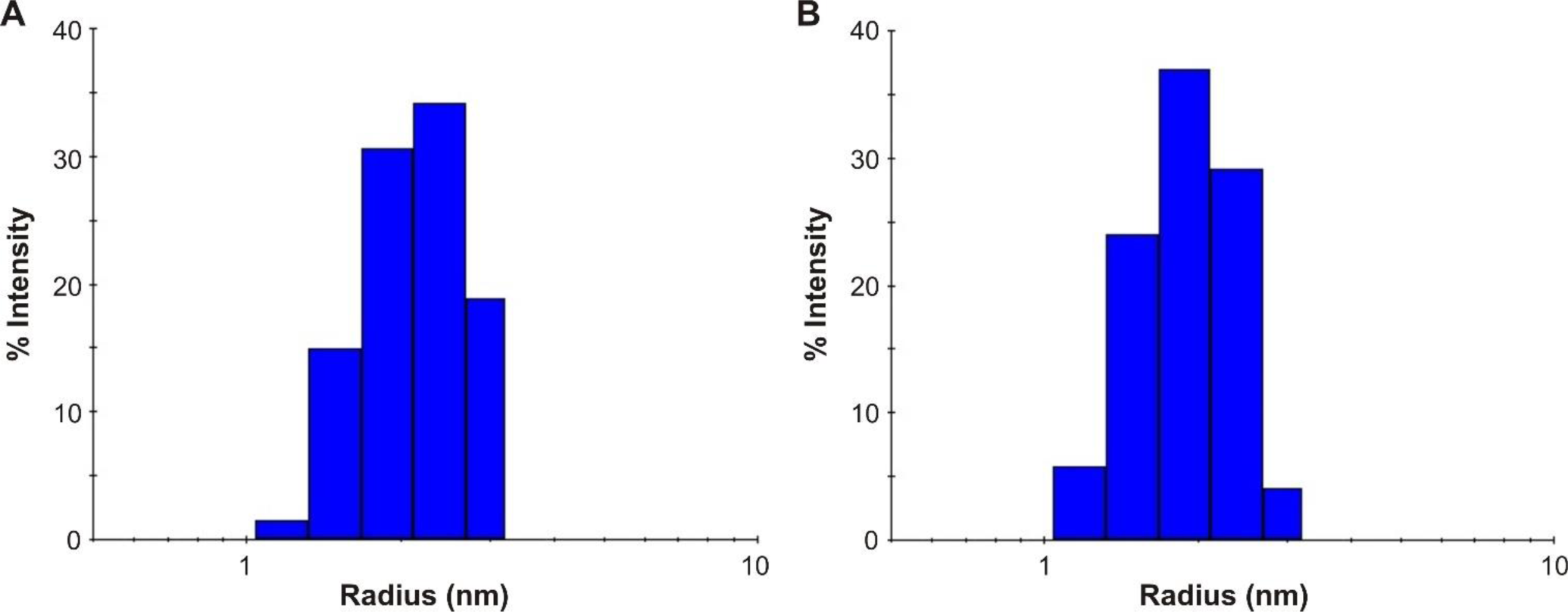
Dynamic light scattering measurements of purified human Atg8-family proteins used in the present reconstitution studies. (**A, B**) Histograms of size distributions of purified LC3B-His12 (**A**) and GATE-16-His12 (**B**) proteins, measured by dynamic light scattering (DLS) using a DynaPro NanoStar DLS instrument (Wyatt Technology). Based upon the measured hydrodynamic radii of human Atg8-family proteins, the molar masses were estimated to be approximately 21 kDa and 16 kDa for LC3B-His12 and GATE-16-His12, respectively.

**Figure S2.**
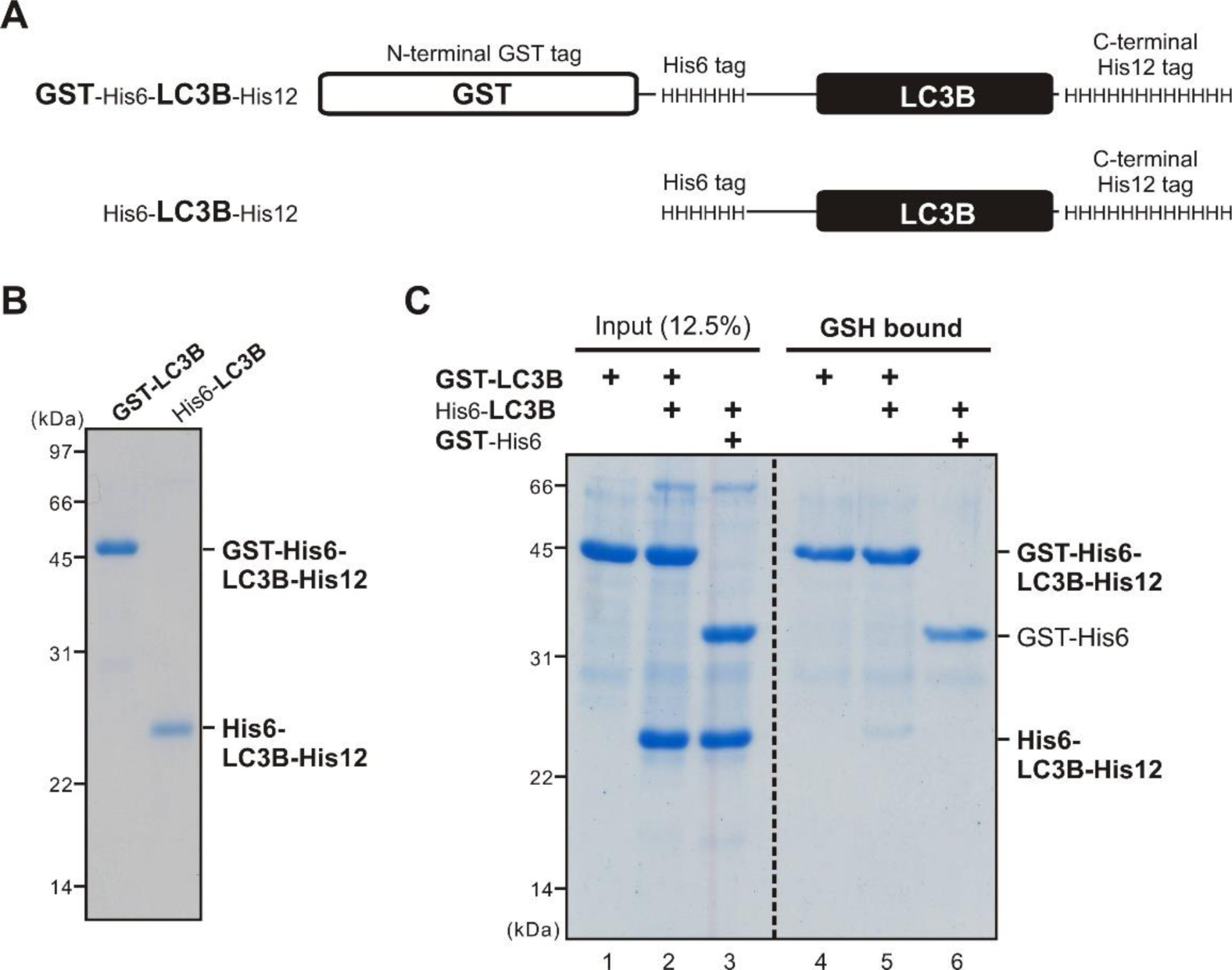
GST pull-down assays to test homo-oligomerization of human LC3B proteins in solution. (**A**) Schematic representation of recombinant LC3B proteins used for GST pull-down assays, GST-His6-LC3B-His12 (GST-LC3B) and His6-LC3B-His12 (His6-LC3B). (**B**) The Coomassie Blue-stained gel of purified GST-LC3B and His6-LC3B proteins used in GST pull-down assays in (**C**). (**C**) Human LC3B has no or little potency to assemble into a homo-oligomer or homo-dimer complex in solution. Purified proteins of GST-LC3B (8 μM final) and His6-LC3B (16 μM final) were mixed in RB150 buffer (20 mM Hepes-NaOH, pH 7.4, 150 mM NaCl, 10% glycerol; 500 μl), incubated with gentle agitation (30°C, 1 h), supplemented with COSMOGEL GST-Accept beads (50% slurry, 200 μl; Nacalai Tesque), and further incubated (4°C, 2 h). After washing the protein-bound GST-Accept beads four times with RB150, GST-LC3B and His6-LC3B proteins bound to the beads were eluted with RB150 containing 100 mM reduced glutathione, and then subjected to SDS-PAGE and Coomassie Blue staining. For the control, purified GST-His6 protein was also incubated with His6-LC3B, mixed with the GST-Accept beads, and analyzed by SDS-PAGE, as employed with GST-LC3B proteins.

**Figure S3.**
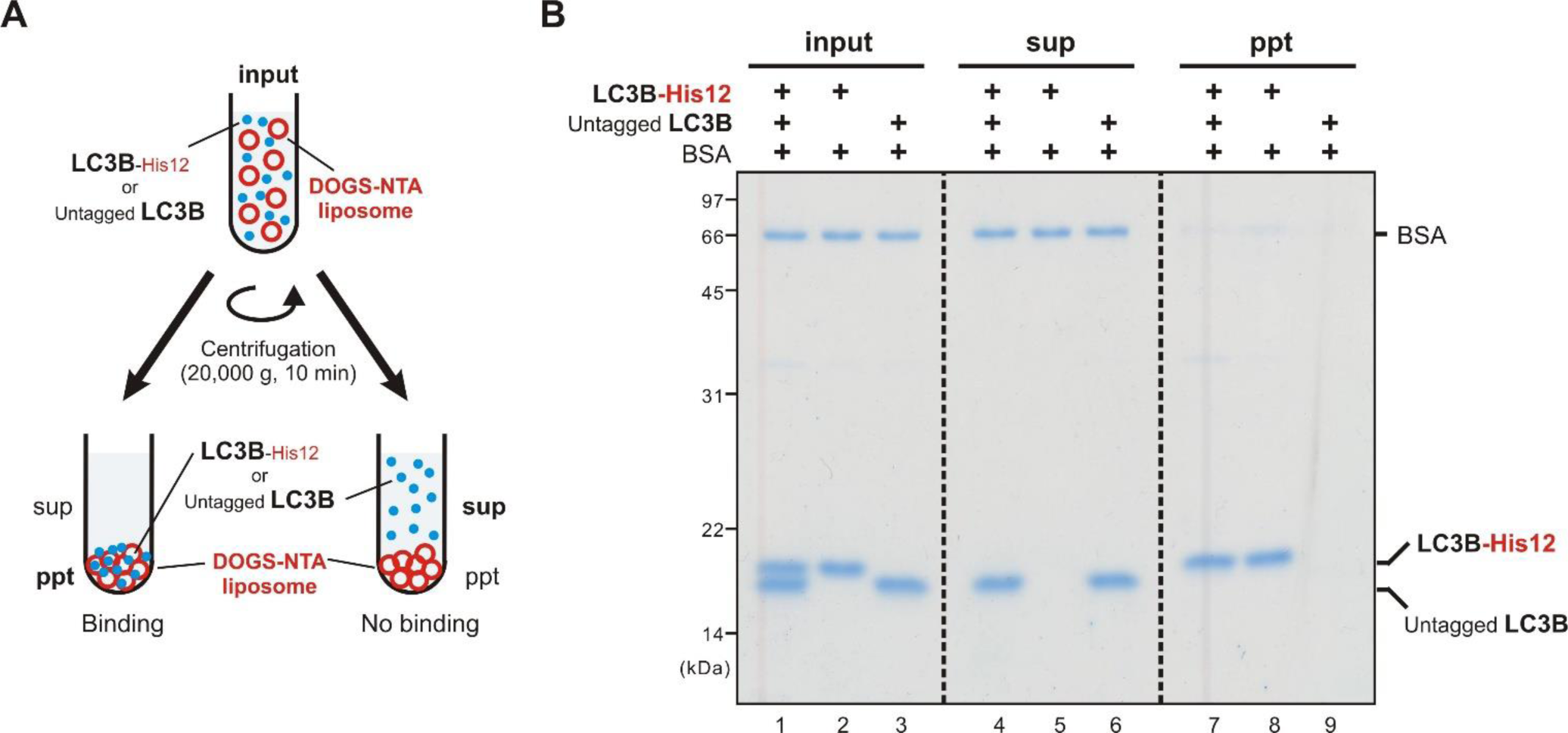
Liposome co-sedimentation assays to test the association of membrane-anchored LC3B proteins with soluble LC3B proteins. (**A**) Schematic representation of liposome co-sedimentation assays testing the interactions of membrane-anchored LC3B-His12 proteins (LC3B-His12) with soluble untagged LC3B proteins lacking a His12 tag (untagged LC3B) on DOGS-NTA-bearing liposomes. (**B**) Untagged LC3B cannot stably associate with membrane-anchored LC3B-His12 on a lipid bilayer. To assay binding of untagged LC3B proteins to LC3B-His12 proteins on liposomal membranes, purified LC3B-His12 (6 μM final), untagged LC3B (6 μM final), and bovine serum albumin (BSA; 3 μM final) were mixed and incubated with extruded liposomes bearing a DOGS-NTA lipid (1.25 mM total lipids in final; 1,000 nm diameter) in RB150. After incubation (30°C, 1 h), the liposome suspensions were centrifuged (20,000 x g, 10 min, 4°C). The input samples before incubation (input) and the precipitates (ppt) and supernatants (sup) obtained after centrifugation were analyzed by SDS-PAGE and Coomassie blue staining.

## References

[1] Klionsky DJ, Ohsumi Y (1999) Vacuolar import of proteins and organelles from the cytoplasm. Annu Rev Cell Dev Biol 15:1–32.

[2] Levine B, Klionsky DJ (2004) Development by self-digestion: molecular mechanisms and biological functions of autophagy. Dev Cell 6:463–477.

[3] Nakatogawa H, Suzuki K, Kamada Y, Ohsumi Y (2009) Dynamics and diversity in autophagy mechanisms: lessons from yeast. Nat Rev Mol Cell Biol 10:458–467.

[4] Mizushima N, Yoshimori T, Ohsumi Y (2011) The role of Atg proteins in autophagosome formation. Annu Rev Cell Dev Biol 27:107–132.

[5] Dikic I, Elazar Z (2018) Mechanism and medical implications of mammalian autophagy. Nat Rev Mol Cell Biol 19:349–364.

[6] Wollert T (2019) Autophagy. Curr Biol 29:R671–R677.

[7] Kriegenburg F, Ungermann C, Reggiori F (2018) Coordination of Autophagosome-Lysosome Fusion by Atg8 Family Members. Curr Biol 28:R512–R518.

[8] Zhao YG, Zhang H (2019) Autophagosome maturation: An epic journey from the ER to lysosomes. J Cell Biol 218:757–770.

[9] Tsukada M, Ohsumi Y (1993) Isolation and characterization of autophagy-defective mutants of Saccharomyces cerevisiae. FEBS Lett 333:169–174.

[10] Klionsky DJ, Cregg JM, Dunn WA Jr, Emr SD, Sakai Y, Sandoval IV, Sibirny A, Subramani S, Thumm M, Veenhuis M, Ohsumi Y (2003) A unified nomenclature for yeast autophagy-related genes. Dev Cell 5:539–545.

[11] Ichimura Y, Kirisako T, Takao T, Satomi Y, Shimonishi Y, Ishihara N, Mizushima N, Tanida I, Kominami E, Ohsumi M, Noda T, Ohsumi Y (2000) A ubiquitin-like system mediates protein lipidation. Nature 408:488–492.

[12] Kirisako T, Baba M, Ishihara N, Miyazawa K, Ohsumi M, Yoshimori T, Noda T, Ohsumi Y (1999) Formation process of autophagosome is traced with Apg8/Aut7p in yeast. J Cell Biol 147:435–446.

[13] Kabeya Y, Mizushima N, Ueno T, Yamamoto A, Kirisako T, Noda T, Kominami E, Ohsumi Y, Yoshimori T (2000) LC3, a mammalian homologue of yeast Apg8p, is localized in autophagosome membranes after processing. EMBO J 19:5720–5728.

[14] Kabeya Y, Mizushima N, Yamamoto A, Oshitani-Okamoto S, Ohsumi Y, Yoshimori T (2004) LC3, GABARAP and GATE16 localize to autophagosomal membrane depending on form-II formation. J Cell Sci 117:2805–2812.

[15] Abeliovich H, Dunn WA Jr, Kim J, Klionsky DJ (2000) Dissection of autophagosome biogenesis into distinct nucleation and expansion steps. J Cell Biol 151:1025–1034.

[16] Xie Z, Nair U, Klionsky DJ (2008) Atg8 controls phagophore expansion during autophagosome formation. Mol Biol Cell 19:3290–3298.

[17] Fujita N, Hayashi-Nishino M, Fukumoto H, Omori H, Yamamoto A, Noda T, Yoshimori T (2008) An Atg4B mutant hampers the lipidation of LC3 paralogues and causes defects in autophagosome closure. Mol Biol Cell 19:4651–4659.

[18] Sou YS, Waguri S, Iwata J, Ueno T, Fujimura T, Hara T, Sawada N, Yamada A, Mizushima N, Uchiyama Y, Kominami E, Tanaka K, Komatsu M (2008) The Atg8 conjugation system is indispensable for proper development of autophagic isolation membranes in mice. Mol Biol Cell 19:4762–4775.

[19] Weidberg H, Shvets E, Shpilka T, Shimron F, Shinder V, Elazar Z (2010) LC3 and GATE-16/GABARAP subfamilies are both essential yet act differently in autophagosome biogenesis. EMBO J 29:1792–1802.

[20] Nakatogawa H, Ichimura Y, Ohsumi Y (2007) Atg8, a ubiquitin-like protein required for autophagosome formation, mediates membrane tethering and hemifusion. Cell 130:165–178.

[21] Weidberg H, Shpilka T, Shvets E, Abada A, Shimron F, Elazar Z (2011) LC3 and GATE-16 N termini mediate membrane fusion processes required for autophagosome biogenesis. Dev Cell 20:444–454.

[22] Nair U, Jotwani A, Geng J, Gammoh N, Richerson D, Yen WL, Griffith J, Nag S, Wang K, Moss T, Baba M, McNew JA, Jiang X, Reggiori F, Melia TJ, Klionsky DJ (2011) SNARE proteins are required for macroautophagy. Cell 146:290–302.

[23] Yang A, Li Y, Pantoom S, Triola G, Wu YW (2013) Semisynthetic lipidated LC3 protein mediates membrane fusion. Chembiochem 14:1296–1300.

[24] Wu F, Watanabe Y, Guo XY, Qi X, Wang P, Zhao HY, Wang Z, Fujioka Y, Zhang H, Ren JQ, Fang TC, Shen YX, Feng W, Hu JJ, Noda NN, Zhang H (2015) Structural Basis of the Differential Function of the Two C. elegans Atg8 Homologs, LGG-1 and LGG-2, in Autophagy. Mol Cell 60:914–929.

[25] Landajuela A, Hervás JH, Antón Z, Montes LR, Gil D, Valle M, Rodriguez JF, Goñi FM, Alonso A (2016) Lipid Geometry and Bilayer Curvature Modulate LC3/GABARAP-Mediated Model Autophagosomal Elongation. Biophys J 110:411–422.

[26] Shpilka T, Weidberg H, Pietrokovski S, Elazar Z (2011) Atg8: an autophagy-related ubiquitin-like protein family. Genome Biol 12:226.

[27] Legesse-Miller A, Sagiv Y, Porat A, Elazar Z (1998) Isolation and characterization of a novel low molecular weight protein involved in intra-Golgi traffic. J Biol Chem 273:3105–3109.

[28] Sagiv Y, Legesse-Miller A, Porat A, Elazar Z (2000) GATE-16, a membrane transport modulator, interacts with NSF and the Golgi v-SNARE GOS-28. EMBO J 19:1494–1504.

[29] Muller JM, Shorter J, Newman R, Deinhardt K, Sagiv Y, Elazar Z, Warren G, Shima DT (2002) Sequential SNARE disassembly and GATE-16-GOS-28 complex assembly mediated by distinct NSF activities drives Golgi membrane fusion. J Cell Biol 157:1161–1173.

[30] van Meer G, Voelker DR, Feigenson GW (2008) Membrane lipids: where they are and how they behave. Nat Rev Mol Cell Biol 9:112–124.

[31] van Meer G, de Kroon AI (2011) Lipid map of the mammalian cell. J Cell Sci 124:5–8.

[32] Vance JE (2015) Phospholipid synthesis and transport in mammalian cells. Traffic 16:1–18.

[33] Yang Y, Lee M, Fairn GD (2018) Phospholipid subcellular localization and dynamics. J Biol Chem 293:6230–6240.

[34] Tamura N, Mima J (2014) Membrane-anchored human Rab GTPases directly mediate membrane tethering in vitro. Biol Open 3:1108–1115.

[35] Inoshita M, Mima J (2017) Human Rab small GTPase- and class V myosin-mediated membrane tethering in a chemically defined reconstitution system. J Biol Chem 292:18500–18517.

[36] Mima J (2018) Reconstitution of membrane tethering mediated by Rab-family small GTPases. Biophys Rev 10:543–549.

[37] Segawa K, Tamura N, Mima J (2019) Homotypic and heterotypic trans-assembly of human Rab-family small GTPases in reconstituted membrane tethering. J Biol Chem 294:7722–7739.

[38] Dennison SM, Bowen ME, Brunger AT, Lentz BR (2006) Neuronal SNAREs do not trigger fusion between synthetic membranes but do promote PEG-mediated membrane fusion. Biophys J 90:1661–1675.

[39] Ji H, Coleman J, Yang R, Melia TJ, Rothman JE, Tareste D (2010) Protein determinants of SNARE-mediated lipid mixing. Biophys J 99:553–560.

[40] Zick M, Stroupe C, Orr A, Douville D, Wickner WT (2014) Membranes linked by trans-SNARE complexes require lipids prone to non-bilayer structure for progression to fusion. Elife 3:e01879.

[41] Hernandez JM, Kreutzberger AJ, Kiessling V, Tamm LK, Jahn R (2014) Variable cooperativity in SNARE-mediated membrane fusion. Proc Natl Acad Sci USA 111:12037–12042.

[42] Geng J, Baba M, Nair U, Klionsky DJ (2008) Quantitative analysis of autophagy-related protein stoichiometry by fluorescence microscopy. J Cell Biol 182:129–140.

[43] Xie Z, Nair U, Geng J, Szefler MB, Rothman ED, Klionsky DJ (2009) Indirect estimation of the area density of Atg8 on the phagophore. Autophagy 5:217–220.

[44] Nagle JF, Tristram-Nagle S (2000) Structure of lipid bilayers. Biochim Biophys Acta 1469:159–195.

[45] Knorr RL, Nakatogawa H, Ohsumi Y, Lipowsky R, Baumgart T, Dimova R (2014) Membrane morphology is actively transformed by covalent binding of the protein Atg8 to PE-lipids. PLoS One 9:e115357.

[46] Nguyen N, Shteyn V, Melia TJ (2017) Sensing Membrane Curvature in Macroautophagy. J Mol Biol 429:457–472.

[47] Erickson HP (2009) Size and shape of protein molecules at the nanometer level determined by sedimentation, gel filtration, and electron microscopy. Biol Proced Online 11:32–51.

[48] Itakura E, Kishi-Itakura C, Mizushima N (2012) The hairpin-type tail-anchored SNARE syntaxin 17 targets to autophagosomes for fusion with endosomes/lysosomes. Cell 151:1256–1269.

[49] Matsui T, Jiang P, Nakano S, Sakamaki Y, Yamamoto H, Mizushima N (2018) Autophagosomal YKT6 is required for fusion with lysosomes independently of syntaxin 17. J Cell Biol 217:2633–2645.

[50] Gao J, Reggiori F, Ungermann C (2018) A novel in vitro assay reveals SNARE topology and the role of Ykt6 in autophagosome fusion with vacuoles. J Cell Biol 217:3670–3682.

[51] Nath S, Dancourt J, Shteyn V, Puente G, Fong WM, Nag S, Bewersdorf J, Yamamoto A, Antonny B, Melia TJ (2014) Lipidation of the LC3/GABARAP family of autophagy proteins relies on a membrane-curvature-sensing domain in Atg3. Nat Cell Biol 16:415–424.

[52] Lo SY, Brett CL, Plemel RL, Vignali M, Fields S, Gonen T, Merz AJ (2012) Intrinsic tethering activity of endosomal Rab proteins. Nat Struct Mol Biol 19:40–47.

